# Maternal exposure to environmental levels of carbamazepine induces mild growth retardation in mouse embryos

**DOI:** 10.1101/2023.01.12.523650

**Authors:** Douek-Maba Orit, Kalev-Altman Rotem, Mordehay Vered, Hayby-Averbuch Hilla, Shlezinger Neta, Chefetz Benny, Sela-Donenfeld Dalit

## Abstract

As chemical pollution is constantly increasing, the impact on the environment and public health must be investigated. This study focuses on the anticonvulsant drug carbamazepine (CBZ), which is ubiquitously present in the environment. Due to its physicochemical properties and stability during wastewater treatment, CBZ is detected in reclaimed wastewater, surface water and groundwater. In water-scarce regions heavily relying on treated wastewater for crop irrigation, CBZ is detected in arable land, produce and even in humans consuming crops irrigated with recealimed wastewater. Aalthough environmental levels of CBZ are very low, risks associated with unintentional exposure to CBZ are essential to be revealed.

In perinatal medicine, CBZ is a teratogen; its prescription to pregnant women increases the risk for fetal malformations. This raises the concern of whether environmental exposure to CBZ may also impact embryogenesis. Studies in zebrafish and chick embryos or in cell culture have indicated negative outcomes upon exposure to low CBZ levels. Yet, these systems do not recapitulate the manner by which human fetuses are exposed to pharmaceuticals via maternal uptake.

Here, we employed the mouse model to determine whether maternal exposure to environmental-relevant doses of CBZ will impact embryonic development. No effects on fertility, number of gestation sacs, gross embryonic malformations or fetal survival were detected. Yet, embryos were growth-delayed compared to controls (*p*=0.0011), as manifested in lower embryonic stage and somite number, earlier morphological features and reduction in mitotically-active cells.

This study provides the first evidence for the effect of environmental concentration of CBZ on the developmental kinetics of maternally-exposed mammalian embryos. While the developmental delay was relatively modest, its consistency in high number of biological replicates, together with the known implication of developmental delay on post-natal health, calls for further in-depth risk analyses to reveal the effects of pharmaceuticals released to the environment on public health.

## INTRODUCTION

Psychoactive pharmaceuticals are among the most frequently-prescribed drugs in the population (Calisto and Esteves, 2009). Many of these drugs find their way to the environment, either via human excretion or by industrial contamination. Among them, the anticonvulsant and mood-stabilizer drug Tegretol (Carbamazepine; CBZ), a sodium channel blocker which modulates neuronal activity, is one of the most persistent pharmaceuticals in effluents (António F. Ambrósio et al., 2002; Leclercq et al., 2009; Post et al., 1983; Qiang et al., 2016; Santos et al., 2007). Due to its long half-life time, physico-chemical properties and ineffective removal in wastewater treatment plants, CBZ and its metabolites reach treated wastewater, and spread into surface water, groundwater and even drinking water (Björlenius et al., 2018a; Brandão et al., 2013a; Hai et al., 2018a; Komesli et al., 2015a; Miao et al., 2005a; Reemtsma et al., 2006a; Verlicchi et al., 2012b). Moreover, CBZ was found to accumulate in the soil and in a variety of crops irrigated with treated wastewater (Beltrán et al., 2020a; ben Mordechay et al., 2018; Malchi et al., 2014a; Paz et al., 2016a), raising the possibility that it enters the food-chain. Notably, in a randomized controlled trial, we have previously found that healthy individuals consuming vegetables irrigated with treated wastewater excrete CBZ and its metabolites in their urine, unlike subjects consuming fresh water-irrigated vegetables (Paltiel et al., 2016a). A more recent follow-up cross-sectional study has investigated the presence of CBZ in the urine of different groups of Israelis, including omnivorous, healthy adults, pregnant women, children, and vegetarians/vegans (Schapira et al., 2020). Detectable levels of CBZ were found in substantial proportion of examined participants (75.9% of adults, 19.6% of children), none of whom were prescribed with the drug. These findings, along with newly published data (ben Mordechay et al., 2022), strongly suggest that people living in regions with widespread irrigation with treated wastewater are unknowingly exposed to CBZ from the environment. Although environmental levels of CBZ in treated wastewater are 10^4^-10^6^ folds lower than clinical doses in the plasma (ng L^−1^ *vs* mg L^−1^) (Bernus et al., 1996; Pomati et al., 2006; Qiang et al., 2016; Verlicchi et al., 2012), the possible impact of involuntary chronic human exposure to environmentally persistent pharmaceuticals must be determined.

In perinatal medicine, CBZ is considered a teratogen (Diav-Citrin et al., 2001; Ornoy et al., 2017). Multiple epidemiological studies have found a higher rate of congenital malformations in fetuses whose mothers were prescribed with CBZ during pregnancy (Barrett and Richens, 2003; Galappatthy et al., 2018; Thomas et al., 2021; Tomson and Battino, 2012; Vajda et al., 2020, 2016). Notably, CBZ can easily pass placental membranes where it can affect placental transport mechanisms as well as reach the fetus due to its neutral and lipid-soluble properties (Kaushik et al., 2016a; Matalon et al., 2002; Tetro et al., 2021). Reported anomalies in the exposed-embryos included neural tube defects as well as malformations in the craniofacial, cardiac, and urinary systems. Fetal growth retardation (FGR) as well as cognitive deficits in the offspring were also reported (Matalon et al., 2002; Neels et al., 2004; Verlicchi et al., 2012). Experiments in rodents have confirmed these findings by showing that intraperitoneal administration of CBZ to pregnant mice at clinically-comparable doses induced malformations (Afshar et al., 2010; Azarbayjani and Danielsson, 1998). Markedly, while these epidemiological and experimental evidences demonstrated the susceptibility of mammalian embryos to clinical doses of CBZ, it is currently not known whether unintentional exposure of healthy pregnant women to CBZ from the environment can potentially risk their fetus. This issue has to be investigated since numerous adult diseases are suspected to have perinatal origin, many of which are still elusive (Nobile et al., 2022).

As experimental exposure of human fetuses to CBZ from the environment is unfeasible, model systems are required. Numerous studies in fish have found that exposure of embryos and larvae to environmental-relevant levels of CBZ via the water-tank led to altered hatching time, reduced body length, small increase in mortality rate as well as modified behavior, implying that teleost embryos are sensitive to environmental-relevant doses of CBZ (Galus et al., 2013a, 2013b; Qiang et al., 2016; Ribbenstedt et al., 2022a; Shao et al., 2019; Zhou et al., 2019). Yet as opposed to avian and mammalian, teleost embryos are not surrounded by amniotic sac. Nonetheless, the impact of environmental levels of CBZ on amniotic embryos is much less known. In a recent study, we examined the effect of environmental-relevant concentrations of CBZ on amniotes, using the chick embryos as an initial proof-of concept model system for embryo-toxicity analysis (Beedie et al., 2016; Davey et al., 2018; Henshel et al., 2002; Kohl et al., 2019). CBZ was added at various environmental concentrations (0.01-100μg L^-1^) on top of the embryo within the egg and found to impair chick embryonic development in a dose- and stage-dependent manner. Effects were exhibited when CBZ was applied at early embryonic stages (gastrulation and neurulation) rather than during later stages of organogenesis. Dose-dependent effects ranged from mild growth-retardation to severe malformations in the neural tube, somites and heart (Kohl et al., 2019).

Although these findings indicated that exposure of chicken embryos to environmentally-relevant levels of CBZ can impair their development, the direct addition of CBZ onto the embryo does not recapitulate the natural exposure route of mammalian fetuses to substances, which can only occur by maternal uptake and transmission. Hence, it remained to be determined whether administration of CBZ at environmental-relevant concentrations to pregnant mammalian females can result in its transmission to the embryos and in compromising their development. Here we addressed for the first time the question of whether and how the development of mouse embryos is affected upon maternal exposure to environmental levels of CBZ. By adding 0.5μg L^-1^ CBZ to the drinking water of female mice *ad libitum* for different time length before and during gestation, we found statistically significant effects on the embryonic stage, as evident by a mild growth retardation in the exposed embryos.

## MATERIALS AND METHODS

### CBZ solution

CBZ (99% purity, Sigma-Aldrich, Rehovot, Israel) was suspended in sterile tap water to prepare a stock solution of 50mg L^-1^. Solution was sonicated for 15 minutes (min) in 40°C, stirred for 16 hours (hr) at room temperature (RT) in a dark bottle, and stored at −20°C before use. Working solution of CBZ (0.5μg L^-1^) was prepared by further dilution in autoclaved-drinking water used in the animal house facility. To verify the precise concentration of CBZ, samples from the working solutions were analyzed by a High-Resolution LC/MS/MS Q-Exactive with a Dionex RSLC system using a Kinetex® 2.6μm EVO C18 100Å, UHPLC column (Phenomenex, US). Mass spectrometer was operated in positive ionization mode and an ESI Full Scan (FS) was used. Ion source parameters were as follows: Spray Voltage (+): 1300.00V; Capillary Temperature: 256°C; Sheath Gas: 51.00; Aux Gas: 3.00; Sweep Gas: 0.00; Probe Heater Temperature: 413°C. Data was analyzed using Tracefinder software. Limit of quantification was for 0.25ng ml^-1^.

### Mice procedures

All procedures were approved by the Hebrew University Animal Care Committee ethical (permit number MD-20-16314-3). Wild-type (WT) male and female C57BL/6J mice were purchased from Harlan Laboratories (Rehovot, Israel). Mice were housed in a specific pathogen-free (SPF) animal house with 12hr light/12h dark cycles (7:00-19:00) with stable temperature and humidity conditions (∼22°C and ∼55%, respectively) and were given autoclaved food and water (Kalev-Altman et al., 2022, 2020). At the onset of sexual maturity (6 weeks), female mice were randomly divided into two experimental groups comprising 20-23 mice each, which received standard drinking water or water containing 0.5μg L^-1^ CBZ *ad libitum*. Water bottles were protected from direct light to avoid <u>photo-degradation</u> of the pharmaceutical, and the level of water was monitored, confirming uniform consumption across the entire experiment. Females were exposed to CBZ for 4-12 weeks before mating. List of all females with their individual age and exposure time to CBZ is provided in Table S1. LC/MS/MS analysis was performed at 8 random time points during the entire experimental period to confirm the CBZ concentration. For mating, each nulliparous female was weighted and housed with a single male for precisely 16h. This time-frame was chosen to reduce the variability in fertilization time between different females. Females were examined for sperm-positive vaginal plug on the following morning, which was considered as embryonic day 0.5 (E0.5). At E9.5, E14.5 and E18.5, females were euthanized and embryos were dissected out of the uterus and washed with phosphate buffered saline (PBS) (Biological Industrial, Beit-Haemek, Israel) for further analyses (Tilleman et al., 2010).

### Analysis of embryonic litter size, morphology and vitality

Gestation sacs were counted in each female to calculate the embryonic litter size and to view vital or absorbed embryos. Embryos were visualized under a stereomicroscope (Discovery V.20 ZEISS, US) to image their size, classify their embryonic stage and count their somites number. Embryos have also been evaluated for different anatomical and morphological parameters to determine the developmental state of particular tissues. Numerical scores were provided for each anatomical parameter, according to subsequent developmental stages of a typical embryo (Table S2). Control or CBZ-exposed embryos were scored for each parameter using a blinded-test. Sum of all criteria resulted in a numerical value of the overall developmental status.

### Flow cytometry

At E9.5, embryos were collected, washed with PBS and dissociated into single cells using collagenase type 4 enzyme at 200Units/ml (Worthington, Lakewood, NJ, US) for 10min at 37°C. Suspended cells were further dissociated by pipetting up and down for 1min, neutralized with PBS and fixated in 4% paraformaldehyde (PFA) (Sigma-Aldrich, Rehovot, Israel) for 2hr at 4°C. Next, cells were centrifuged at 13000g four times, 5min each and washed in PBS in-between each centrifugation. Cells were then incubated with a blocking solution (0.2% Triton in PBS and 5% goat serum) for 2hr at RT, followed by an overnight incubation at 4°C in blocking solution with anti phospho-histone 3 (PhH3) antibody or anti-caspase 3 (Cas3) antibody (both 1:300; Lifespan Bioscience. Washington US). Next, cells were re-centrifuged and washed in PBS four times before incubated for 2hr at RT with Alexa-Fluor 488 goat anti-rabbit antibody (1:300; Molecular probes, Oregon, USA). Cells were re-centrifuged, and washed with PBS three times. Finally, PBS-suspended cells were passed through a cell-strainer cap (Life Science, Leipzig, Germany) for separation into single cells, then run through an Accuri C6 Flow Cytometer (BD Biosciences, CA, USA). Flow cytometry analysis was performed using BD Accuri C6 software as was previously described (Kohl et al., 2019; Peretz et al., 2016).

### Data analysis and statistics

All data are expressed as the mean ± SD (standard deviation). The significance of differences between groups was determined using JMP 15.0 Statistical Discovery Software (SAS Institute 2000) and GraphPad Prism 8 software (GraphPad Software, CA, US) by one-way ANOVA followed by a Tukey Kramer HSD test, unpaired T-test with Mann Whitney test or Fisher exact test. When indicated, dot plots with color scales were created in-house in R using the ggplot2 package (Table S3). Differences were considered significant at P<0.05. Groups with the same letter were not measurably different, and groups that were measurably different are indicated by different letters. Groups may have more than one letter to reflect the “overlap” between the sets of groups, and sometimes a set of groups is associated with only a single treatment level.

## RESULTS AND DISCUSSION

### Exposure of female mice to an environmental-relevant level of CBZ

The frequent detection of CBZ in various aquatic environments (Leclercq et al., 2009; Qiang et al., 2016; Wilkinson et al., 2022), together with its teratogenic activity (Barrett and Richens, 2003; Ferrari et al., 2003; Matalon et al., 2002), calls for assessing whether environmental-relevant concentrations of CBZ can impair embryonic development. Notably, negative effects were previously reported in chick and fish embryos upon their exposure to environmental levels of CBZ through the egg or water tank (Galus et al., 2013a, 2013b; Hermsen et al., 2013, 2011; Kaushik and Thomas, 2019; Kohl et al., 2019; Pohl et al., 2021; Ribbenstedt et al., 2022; Zhou et al., 2019). Yet, as these model systems do not recapitulate the exposure route of human fetuses to such contaminants, prediction of potential risks to humans cannot be fully anticipated by these studies. Here, we have utilized mouse embryos as the gold-standard mammalian model system that recapitulates human development and disease, to investigate whether embryonic development is compromised upon exposure of female mice to environmental-relevant levels of CBZ. Sexually matured (6 weeks old) nulliparous females were given drinking water supplemented with 0.5μg L^-1^ CBZ. This concertation represents CBZ levels reported in treated wastewater, groundwater and surface water around the world (Hai et al., 2018; Paltiel et al., 2016b; Wilkinson et al., 2022), and hence reflects a relevant scenario of unintentional exposure to CBZ from the environment. In parallel, age-matched control females received drinking water with no additional drug. Water-bottles were randomly tested for CBZ concentration, revealing an average concentration was 0.495μg L^-1^ (median of 0.435μg L^-1^; max 0.665μg L^-1^; min 0.31μg L^-1^). As the average daily consumption of water for a 25gr adult mouse is ∼4 ml (Bachmanov et al., 2002), each mouse was exposed to ∼0.08 ng/g body weight (2 ng/mouse) per day. Females were provided with CBZ for variable time lengths before gestation (30-84 days), in order to test whether different exposure times to CBZ correlate with specific embryonic phenotypes. The minimal exposure period of 30 days was selected based on a previous study which showed that exposure of female mice to 100μg L^-1^ CBZ for 20 days (10 days pre- and 10 days post-fertilization) was sufficient for the transmission of CBZ to their embryonic tissues, such as liver and brain (Kaushik et al., 2016a). All females were housed with males for the same time window (16hr), in order to obtain timed pregnancies in individual females. Uteri were extracted at gestation day (GD) 9.5, 14.5, and 18.5 for analyses at different embryonic stages. A schematic illustration of the experimental plan is presented in Fig. 1.

**Figure 1.**
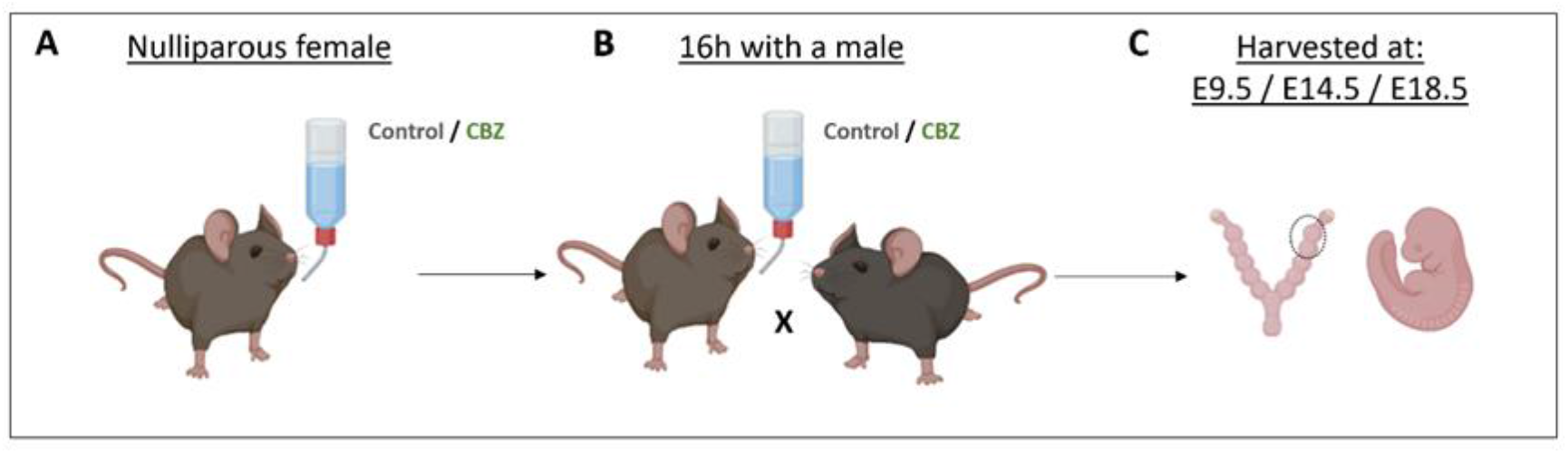
Experimental design. (A) Nulliparous females were exposed through their drinking water to 0.5μg L^-1^ CBZ and compared to control mice receiving standard drinking water. (B) Females were paired with males to become pregnant. (C) Females were scarified at gestation day 9.5, 14.5, or 18.5. Uteri were harvested and counted for the number of successfully implanted gestation sacs or resorbed sacs as well as for embryonic vitality and morphology. Illustration was created with BioRender.com.

### Maternal exposure to CBZ has no effect on embryonic implantation, litter size or viability

To evaluate whether exposure of females to environmental level of CBZ affects their fertility, number of gestation sacs as well as embryonic survival were assessed. Uteri from control (n=20) and CBZ-treated females (n=23) were analyzed for the presence of gestation sacs at GD9.5, GD14.5 and GD18.5. Number of sacs varied and ranged between 2-13, with an average of ∼7 gestation sacs in both control or CBZ-treated groups, with no statistically significant differences between the groups (Fig. 2A). Additionally, no significant differences were found between the control and CBZ groups in the mean number of viable embryos (∼7) or in the number of resorbed sacs (∼1), which reflected successful implantations with no succeeding embryonic mortality (Fig. 2 B, C). Notably, although individual females varied in the numbers of normal *vs* resorbed sacs, these variations were found in both groups, with no association to CBZ exposure (Fig. 2 D-D’).

**Figure 2.**
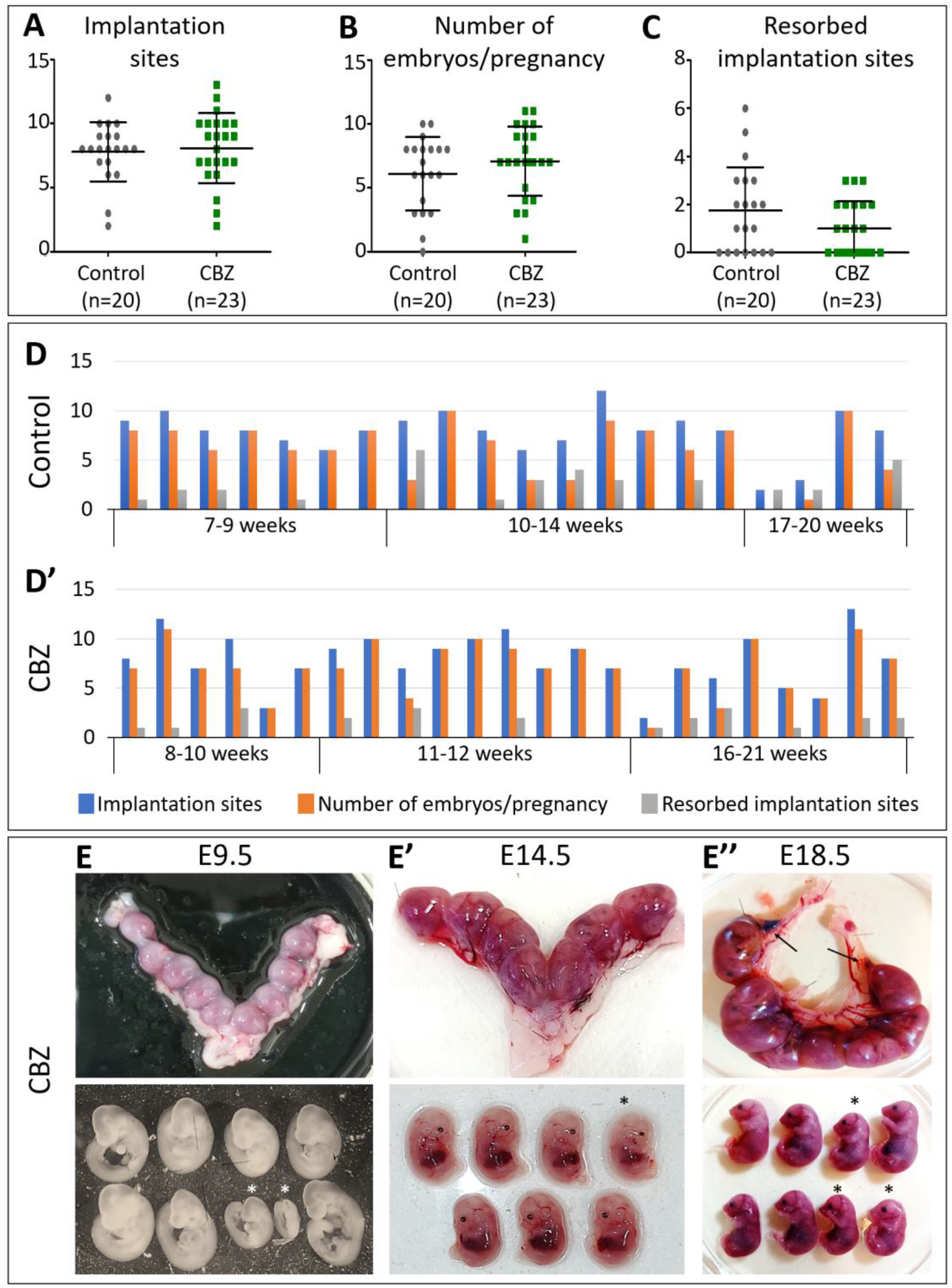
The effect of CBZ on implantation, litter-size and embryonic mortality. (A) Total number of implantation sites in uteri collected from control or CBZ-exposed females (ns, P=0.65). (B) Number of resorbed sacs per pregnancy of control or CBZ-exposed female (ns, P=0.19). (C) Number of sacs with viable embryos per pregnancy of control or CBZ-exposed female (ns, P=0.29). Each dot in (A-C) represents the average value of all embryos within one female. (D-D’) Individual variation in the measured parameters in both groups with no significant differences between control and CBZ-treated mice. (E-E’’) Representative uteri at GD9.5, GD14.5 and GD18.5 from CBZ-treated females including some resorbed sacs (E’’, arrows) and the harvested embryos from each. While most embryos of the same litter are comparable in their gross size, some seem smaller than others (bottom panel, asterisks). E, embryonic day; CBZ; carbamazepine.

As different females were exposed to CBZ for diverse time lengths, and hence were of different age when mated, we next asked whether the number of sacs with vital embryos correlated with different exposure times to CBZ. In this analysis, females were divided into three groups according to their exposure time to CBZ in weeks (w) (details regarding each female are provided in Table S1). Examination of individual females of both groups revealed some variations in the number of gestation sacs and in the proportion of live/absorbed embryos, but these differences did not correlate with the female’s age and exposure time to CBZ (Fig. 2 D-D’). These results demonstrate that exposure of female mice to low environmental-relevant concentration of CBZ does not impair their ability to conceive, to succeed in embryonic implantation, and to support embryonic survival during pregnancy.

Visualization of the analyzed properties demonstrates typical uteri from control or CBZ-treated females at GD9.5, GD14.5 and GD18.5, which display 9, 7 or 10 embryonic sacs, respectively (Fig. 2 E-E’’, top). Notably, out of the 10 sacs observed at GD18.5, two sacs were resorbed (Fig. 2E’’, arrows). Morphological examination of the embryos did not reveal any gross defect in the CBZ-exposed group in comparison to controls (Fig.2 E-E’’, bottom). Yet, while most embryos within the same uterus were matched in their developmental stage, some embryos seemed smaller (Fig. 2 E-E’’, asterisks). Notably, deviations in the anticipated embryonic stage are known to occur within littermates, due to variable time of conception, implantation time and nutrient disparity (Croy et al., 2014; Kalev-Altman et al., 2020; Wong et al., 2015). However, variations in the developmental kinetics of individual mouse embryos within the same mother were also reported to occur due to maternal exposure to different environmental pollutants (Althali et al., 2019; Guo et al., 2019; Müller et al., 2018; Tao et al., 2022). Hence, it was necessary to further determine whether the observed growth differences were random or associated with maternal exposure to CBZ.

### The association between maternal exposure to CBZ and developmental delay

To examine whether maternal exposure to environmental levels of CBZ is linked to a slower embryonic development, embryos from control and CBZ-exposed mothers at GD9.5 were analyzed in a blinded test for their precise stage based on the classical Theiler staging criteria (Theiler, 1989). Females who had only one live embryo or did not conceive were excluded from the analysis, resulting in an evaluation of 11 control females and 9 CBZ-exposed females, with a total of 72 and 77 embryos in each group, respectively. Our results show that the average stage of control embryos was E9.5, as expected based on the female’s GD. Yet, the average stage of the CBZ-exposed embryos was found to be slightly younger, E9.2 (Fig. 3 A). This difference was found to be statistically significant (P=0.0011), in spite of the fact that the evident embryonic stages at both groups at the harvesting varied (between E8.0-E10.5). To further verify this result, number of somites was counted. Somites begin to appear at E7.5 and increase to a final total of 60 somites at E14.0. Somite number is a quantifiable landmark to determine developmental progression predominantly between E8.0 and E10.5, because there is an increase of 22 somites which are clearly visible in that time frame (Wong et al., 2015). Although the expected somite number at E9.5 can vary between 22-28 pairs (Theiler, 1989), a small but significant difference was found in somite number between the two groups; embryos from CBZ-exposed mothers had a mean of 21.4 somites while the mean number of control embryos was 23.6 somites (Fig. 3 B, P=0.002). Interestingly, this analysis also showed that control embryos varied more in their stage and somite number, in comparison to CBZ-exposed embryos, which displayed more values and hence were more clustered (Fig. 3 A, B). The evident reduction in the developmental stage and somite number did not correlate with the female’s age or the exposure time to CBZ (Fig. 3 C-D’), as females exposed to CBZ for different time lengths gave rise to embryos with similar variations in stage and somite number as control females, with no increase in the occurrence of developmental delay upon longer exposure to CBZ. Together, these results indicate that exposure of female mice to 0.5μg L^-1^ CBZ in their drinking water, leads to a moderate but significant delay in the developmental kinetics of the embryos, as found by a higher proportion of embryos with 10% reduction in their somite number and with a younger-than anticipated embryonic stage at GD9.5.

**Figure 3.**
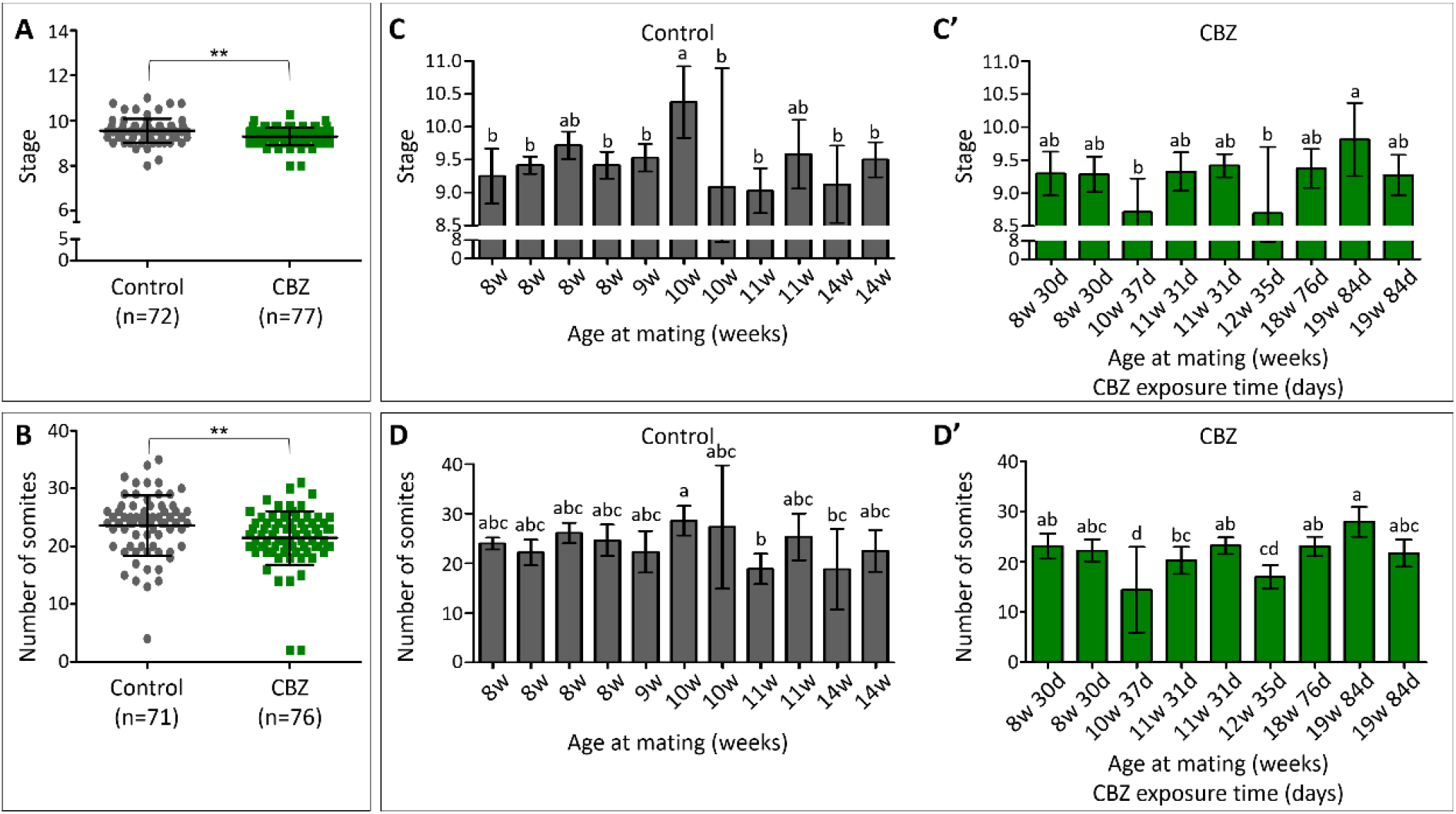
Decreased embryonic stage and reduced somite number upon maternal exposure to CBZ. (A, B) Assessment of embryonic stage and somite number in individual embryos collected from control or CBZ-exposed mothers at GD9.5, using Theiler staging and somite counting. Each dot represents an embryo. The CBZ-group showed a statistically significant decrease in both parameters compared to control (** p< 0.005). (C-D’) Assessment of embryonic stages (C, C’) or somite number (D, D’) in embryos of individual females from the control or CBZ-exposed groups. Each bar represents a mean value of the measured parameter in all embryos from one female, named based on her age in weeks when mated and her exposure time to CBZ in days. Variability in the embryonic stage or number of somites is evident between different females from each group, with no statistically significant differences between different females. Different letters denote significant differences at P<0.05 between groups. CBZ, carbamazepine; w, weeks; d, days.

### Morphological evaluation and scoring of embryos

To further validate the effect of maternal exposure to environmental level of CBZ on embryonic development, we implemented an unbiased scoring method to quantify anatomical differences between embryos. This approach is based on various morphological landmarks that define different tissues at different stages, from E7.5 to E11.25. Scoring was performed in a blinded manner on the following tissues: branchial arches, optic system, heart, neural tube, limbs, and tail (Jacobson and Tam, 1982; Nandi and Mishra, 2015; Papaioannou and Behringer, 2005; van Maele-Fabry et al., 1990; Yang et al., 2010). The morphological state of a certain tissue gained a numerical score which increased according to progression in development (Table S2). Scores of individual features were summed up to provide a final score for the developmental state of each embryo. Slightly lower scores were given to the CBZ group in most of the examined parameters (i.e., branchial arches, eye, heart, forelimb bud,) as compared to the control group (Fig. 4 A), although only the score differences in the branchial arches and eye morphology were found to be statistically significant (P<0.05, P<0.005, respectively) (P values for heart and forelimb bud differences were 0.64 and 0.1, respectively). For instance, many of the CBZ-exposed embryos showed elongated optic primordium with an ovoid shape while most of control embryos showed a more advanced primary optic vesicle with an open optic stalk. Summing up the tissue-individual scores to a global score provided a significant lower value for the CBZ-group compared to the control group (15.48 *vs* 16.06, respectively, P<0.05) (Fig. 4 B). As before, comparison between embryos from different females revealed no correlation between lower or higher scores and the female’s age and exposure-time to CBZ (Fig. 4 C, C’).

**Figure 4.**
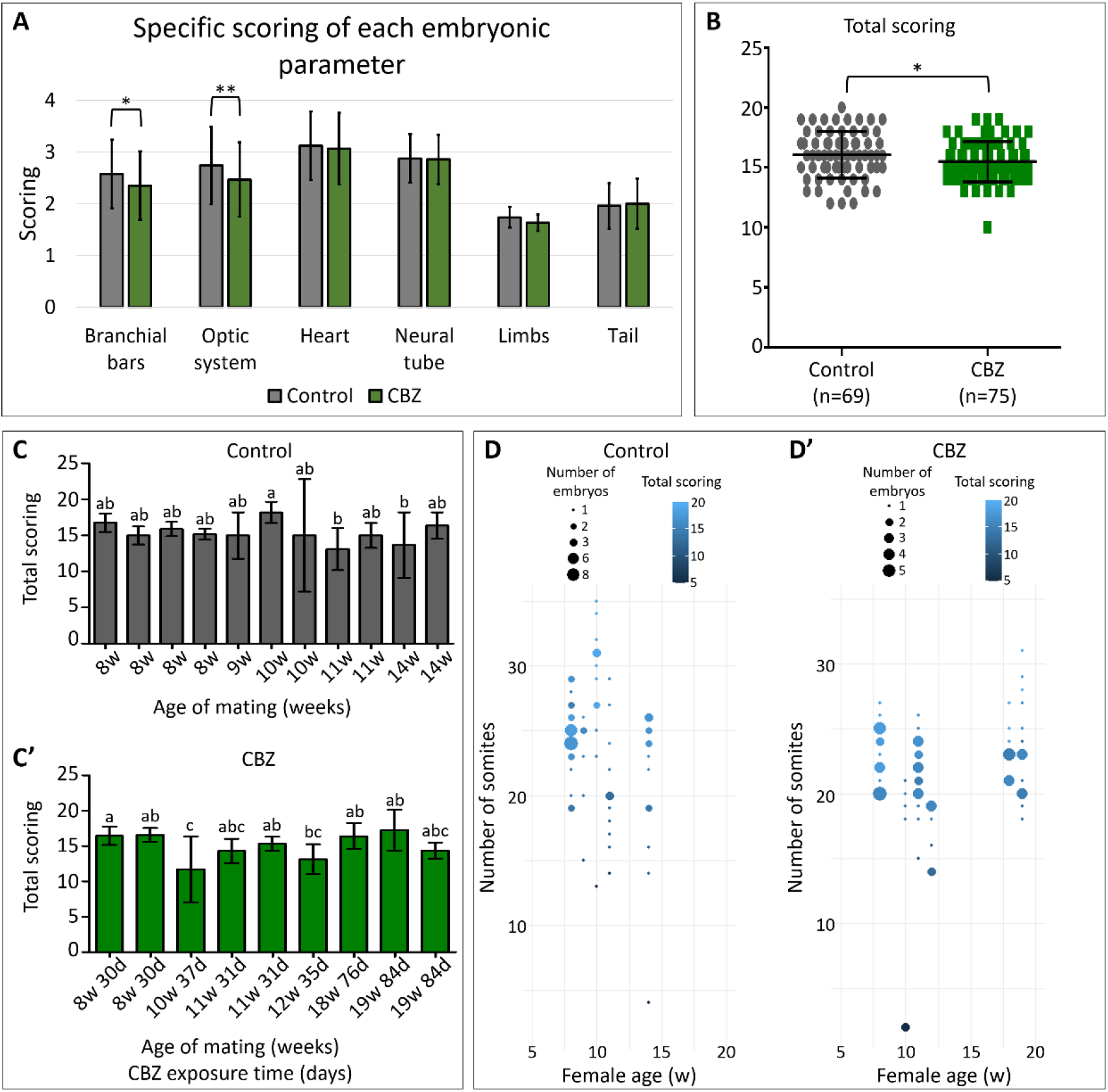
Quantification of the developmental state of control and CBZ-embryos based on morphological criteria. (A) All embryos from each group were analyzed for different morphological parameters at different tissues and were provided with scores that represented the developmental state of each tissue (see also sup Fig. 3). Statistically-significant lower scores were given for branchial arches (* p<0.05) and eye development (** P<0.005) in the CBZ embryonic group compared to controls. Each bar represents a mean value of the measured parameter in all embryos. (B) The global score of all measurements together found to be significantly lower in the CBZ-group compared to control embryos (* p < 0.05). Each dot represents the global score of each embryo. (C, C’) The averaged global morphological score as given to all embryos of each control and CBZ-exposed female. Different letters denote significant differences at P<0.05 between groups. Each bar represents a mean value of the measured parameter in all embryos from one female, named based on her age in weeks when mated (w) and her exposure time to CBZ in days (d). (D, D’) Dot-plot analysis to demonstrate the scattering of embryos from all control and CBZ-exposed females. Each dot represents one or more embryos with similar number of somites together with similar total score. The intensity of the color correlates with the value of the score. The size of each dot represents the group size, namely the number of embryos in each group (see also Table S3).

Importantly, low morphological scores can either represent the morphological state of younger embryos or can result from embryos that develop to the expected age but carry morphological defects. To fully dissociate between the two options, we had to compare the somite stages of individual embryos from all mothers with their morphological score. Distribution of all embryos per all females was analyzed by generating a dot-plot of different embryonic group sizes which cluster based on a similar somite number and morphological score (Fig.4 D, D’, Table S3). Distribution of all embryos shows a clear correlation between the two tested parameters, namely, the lower the somite number is, the lower the morphological score is, whereas the increase in somite number correlates with a gradual increase in embryonic scoring. Moreover, this data provides an overview of the distribution of all embryos in the control and CBZ groups, indicating that a majority of CBZ-exposed embryos displays a combination of fewer somites with lower scores, in comparison to the control ones. This analysis further confirms that the CBZ-exposed embryos are younger-than expected, as indicated by the match of their somite number and morphological score, but do not present malformations in different tissues.

Altogether, these results suggest that exposure of female mice to CBZ at a low environmental concertation leads to a higher proportion of slightly-growth delayed embryos. Yet, this phenotype was not coupled with any apparent malformation in any of the examined tissues, indicating the less-likelihood of CBZ to cause teratogenic effects in mammalian embryos, as opposed to its effect when applied at clinical doses (Barrett and Richens, 2003; Beutler et al., 2005). Interestingly, a recent toxico-metabolomic study in zebrafish which were exposed to varied concentrations of CBZ, reveled a significant delay in hatching upon exposure to a similar lower-dose of CBZ as used in our study (Ribbenstedt et al., 2022), which may suggest a comparable slower developmental kinetics in both species.

Markedly, the mild growth delay found in the mice embryos differed from the phenotypes we previously detected in chick embryos that have been exposed to a similar CBZ concentration. In the chick embryos, a more profound developmental delay and various types of malformations appeared following application of CBZ into the eggs (Kohl et al., 2019). The discrepancy in the CBZ effects between chick and mouse embryos can be attributed to two main scenarios (which are not mutually exclusive): one possibility is that, as opposed to the maternal route of exposure to CBZ in mice embryos, the more-interventional application of CBZ directly on top of the chick embryo makes it less resilient, due to the unavoidable sensitivity to any direct manipulation at early embryonic stages, before organogenesis commenced. The second possibility is that the severer phenotypes in the chick correlate with higher concentration of CBZ that reach the embryos upon the direct application, as opposed to the non-direct CBZ transmission *via* the mother. Markedly, dose-dependent effects were also reported in zebrafish, in which chronic exposure to environmentally relevant levels of CBZ impacted multiple organs, leading to dose-dependent decline in reproduction, increase in organ pathologies, appearance of edemas and mortality (Galus et al., 2013a, 2013b; Pohl et al., 2021). Behavioral tests also demonstrated altered responses of fish to environmental stimuli that were correlated with a gradual increase in environmental CBZ dosages (Qiang et al., 2016; Thomas et al., 2012). Interestingly, while in our study we intentionally used a low level of CBZ to recapitulate the common levels found in groundwater/surface water/treated wastewater, further studies are needed to assess whether a severer growth delay will be found in mouse embryos upon maternal exposure to higher environmental-relevant doses of CBZ, such as those used in previous studies in fish, mouse and in cultured cells in vitro (Kaushik et al., 2016b; Thomas et al., 2012; Zhou et al., 2019).

### CBZ effect on cell proliferation

One mechanism by which CBZ may delay developmental kinetics is by affecting cell division. We have previously found that addition of CBZ at environmental-relevant concentrations to chicken embryos resulted in a marked decrease in mitotically-active cells without increasing cell death (Kohl et al., 2019). Similarly, treatment of rats with clinical doses of oxcarbazepine, a derivative of CBZ, induced cell cycle arrest in renal epithelial cells (Ota et al., 2021). To test whether this phenomenon occurs also in our system, embryos from control (n=11) or CBZ-exposed females (n= 13) at GD9.5 were dissociated into single cells and immunostained for phospho-histone H3 (PhH3), a nuclear marker that specifically labels dividing cells during metaphase (Peretz et al., 2018; Pokhrel et al., 2022; Sawicka and Seiser, 2012). Cells were analyzed by flow cytometry assay to quantify the proportion of PhH3-stained cells out of the total cells (Fig. 5 A). The percentage of PhH3 expressing cells was found to be significantly lower (p<0.005) in the CBZ-embryonic cell group compared to control-embryonic cells (Fig. 5 B, B’, red histogram/12.7% or green histogram/23.7%, respectively). Importantly, this effect was not coupled with an increase in cell death, as evaluated by flow cytometry analysis on cells stained for the apoptotic marker Caspase3 (Crowley and Waterhouse, 2016) in the CBZ-treated group in comparison to the control group (Fig. S1). Together, these results suggest that the developmental delay triggered by maternal exposure to a low environmental concentration of CBZ is associated with a decrease in the number of actively dividing cells in the exposed embryos.

**Figure 5.**
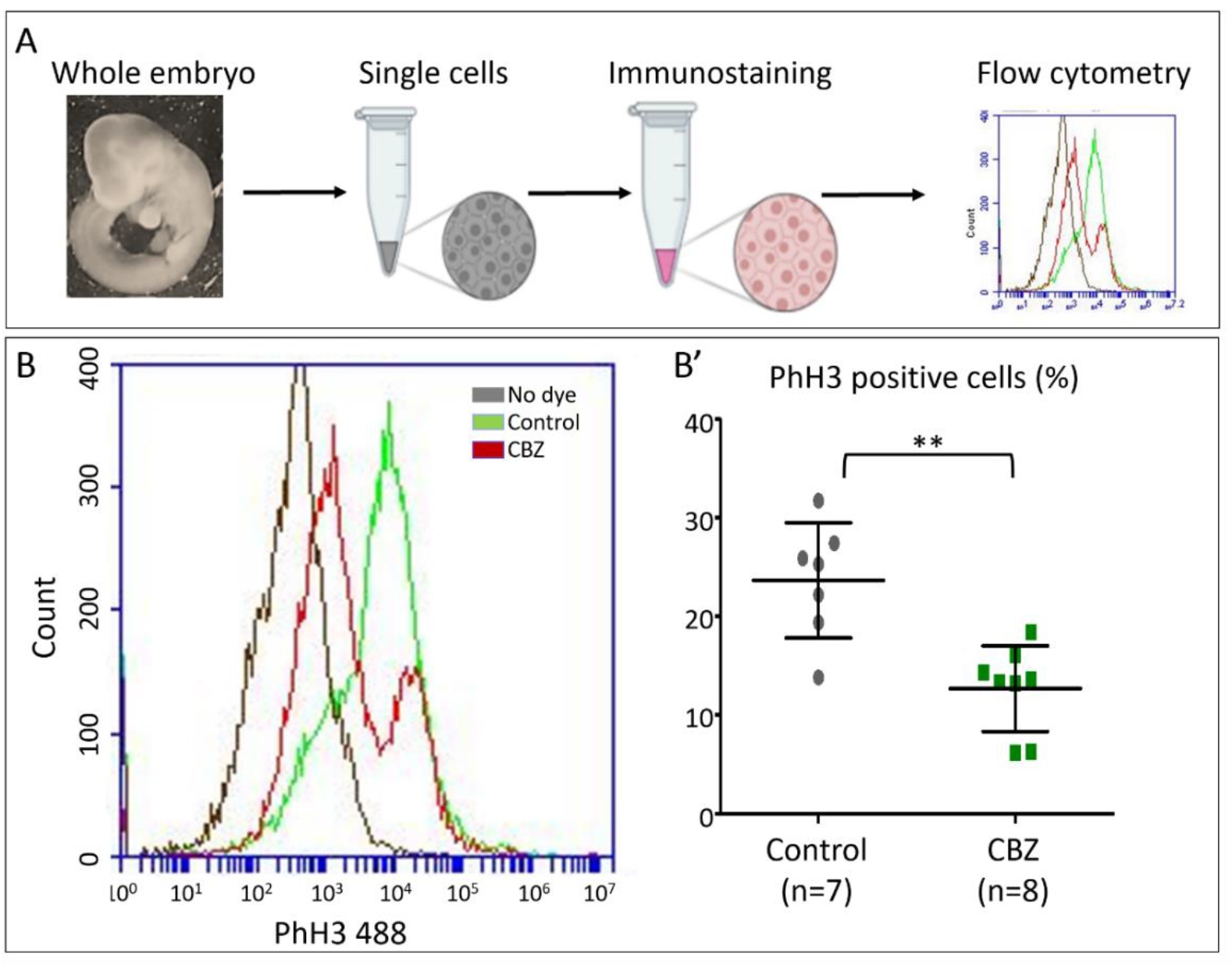
The effect of CBZ on cell proliferation. (A) Schematic illustration of the experimental procedure in which whole embryos were dissociated into single cells, immune-stained with PhH3 antibody and analyzed by flow cytometry. (B) Example histograms for flow cytometric analysis of PhH3-stained cells from control embryos (green histogram) or CBZ-exposed embryos (red histogram). Negative control of on-stained cells is presented in the black histogram. (B’) Quantification of the percentage of PhH3-expressing cells out of the total cells in each group (** p<0.005) from different biological repeats. Each dot represents total cells from a pool of 2-3 different embryos from 4 control females and 6 CBZ-exposed females.

The reduction in the number of mitotically-active cells is in agreement with our previous results in chick embryos (Kohl et al., 2019). Other studies in a human bronchial epithelial cell line further supported our data by showing that alterations in cell-cycle genes (Song et al., 2011), inhibition of hormonal secretion in mammalian (Ahmed et al., 2008), or impairment in centrosome separation in human kidney epithelial cell line (Pérez Martín et al., 2008), were all detected following CBZ treatment, although in these studies clinical or sub-clinical doses of CBZ have been used. Moreover, experiments in rat and zebrafish embryos, as well as in several cell lines of human, primates and murine origin, described similar growth-retardation and/or anti-proliferating effects of CBZ (Ahmed and El-Gareib, 2017; Jos et al., 2003; Pérez Martín et al., 2008; Shaikh Qureshi et al., 2014; Tittle and Schaumann, 1992; van den Brandhof and Montforts, 2010; Weigt et al., 2011). For instance, fish embryos exposed to low CBZ levels (0.5μg L^-1^) exhibited altered hatching time and body length, as well as a small increase in mortality rate and modified behavior (Galus et al., 2013a, 2013b; Qiang et al., 2016). Despite ours and other’s evidences, the molecular mechanism by which CBZ affects cell division remains elusive.

### Possible implications on human health

Our data show a mild but consistent growth delay in embryos obtained from CBZ-exposed mothers at GD9.5 in comparison to controls. This phenomenon was coupled with a decrease in cells at metaphase. It remains to be clarified whether these findings augur a higher frequency of dysmorphogenetic appearance at birth, since even mild perturbations in early embryonic life may forecaster many types of diseases in adults (Nobile et al., 2022). In relevance to our findings, multiple epidemiological studies in humans have shown that fetal growth restriction (FGR), defined as any impairment in fetal growth compared to the expected biological potential in utero, is in association with reduced postnatal survival rates and a greater risk of perinatal morbidity and mortality (Lefebvre et al., 1998; Malhotra et al., 2019; Nardozza et al., 2017). Early FGR also predisposes the development of major malformations, such as neural tube defects, impaired neurological and cognitive development, as well as cardiovascular, respiratory, renal, endocrine and other chronic diseases in children and adults (Spiers, 1982) (Mericq et al., 2017; Chatmethakul and Roghair, 2019). Importantly, FGR in human fetuses has been associated with maternal exposure to environmental pollutants (Gómez-Roig et al., 2021). For instance, maternal exposure to airborne pollutants, mycotoxins, phthalates, as well as to drinking water contaminated with nitrate and pesticide, has been linked to a higher occurrence of FGR (Alvito and Pereira-da-Silva, 2022; Bach et al., 2015; Chang et al., 2022; Gönenç et al., 2020; Macchi et al., 2021; Santos et al., 2021; Smith et al., 2017)As our study demonstrates for the first time a small but consistent growth delay in mice embryos upon maternal exposure to an environmental level of CBZ, further studies are needed to determine whether the mild growth restriction found in the CBZ-exposed mice at GD9.5 persists throughout gestation, in order to conclude whether CBZ from the environment may induce FGR in mice, which may be comparable to clinical evidences in humans (Dilworth et al., 2011). If so, it would appear necessary to begin exploring whether higher incidences of FGR are detected in regions with relatively high levels of CBZ in treated wastewater, surface water and groundwater.

## Conclusion

This study provided the first evidence on a causal link between chronic maternal exposure to an environmental concentration of CBZ and growth delay in mouse embryos. The developmental delay phenotype was not accompanied with noticeable birth-defects or increased embryonic mortality. As developmental delay in human fetuses is associated with compromised development and morbidity which can be extended to adulthood, our data calls for further analysis to decipher whether, and by which mechanisms, the presence of pharmaceuticals in the environment can impair embryonic health and growth.

