## supplemental data for "Maternal exposure to environmental levels of carbamazepine induces mild growth retardation in mouse embryos"

**Supplementary Table 1**: Presentation of all control and CBZ-exposed females used for the study. Females' age at the time of mating is shown in weeks and the exposure time to CBZ is shown in days.

| Control | CBZ | |
| --- | --- | --- |
| female age (weeks) | female age (weeks) | exposure time (days) |
| 7 | 8 | 30 |
| 8 | 8 | 30 |
| 8 | 8 | 30 |
| 8 | 11 | 31 |
| 8 | 11 | 31 |
| 9 | 10 | 37 |
| 9 | 11 | 18 |
| 10 | 11 | 30 |
| 10 | 11 | 30 |
| 10 | 11 | 31 |
| 10 | 11 | 31 |
| 11 | 12 | 35 |
| 11 | 12 | 35 |
| 11 | 12 | 35 |
| 14 | 12 | 35 |
| 14 | 16 | 63 |
| 17 | 17 | 90 |
| 17 | 18 | 24 |
| 20 | 18 | 76 |
| 20 | 18 | 90 |
|  | 19 | 84 |
|  | 19 | 84 |
|  | 21 | 84 |

**Supplementary Table 2:** A numerical scoring method to quantify the morphological state of different embryonic tissues.

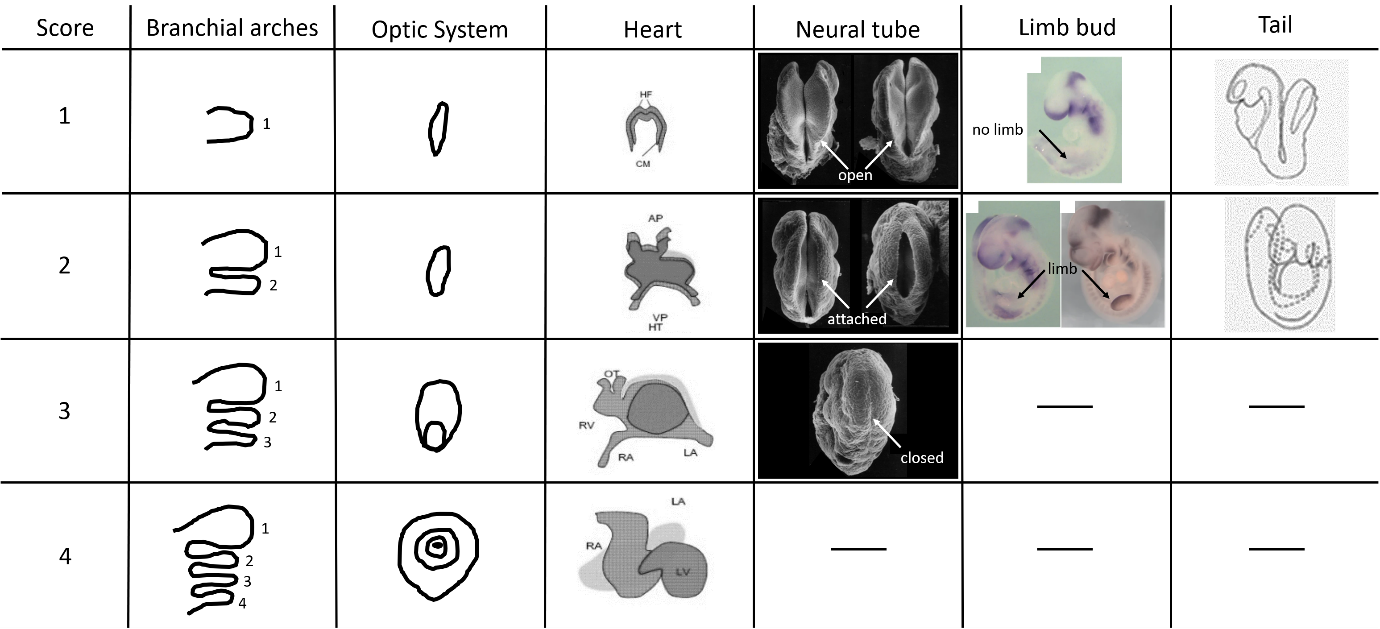

Scores (left column) are based on the following morphological features (left to right): (i) number of branchial arches (1-4); (ii) Optic system- absent eye, elongated optic primordium, ovoid vesicle when an open optic stalk, faint gray hue pigment (1-4); (iii) Heart- primary heart crescents, heart tube formation, heart looping, heart chamber formation (1-4); (iv) Neural tube- open neural tube along its entire axis, partially closed neural tube with open neuropore, closed neural tube (1-3); (v) Forelimb bud- no apparent limb bud, limb bud is present (1,2); (vi) Tail shape- Tail turning outwards. Tail turning inwards (1,2). (Adopted from Jacobson and Tam, 1982; Van Maele-Fabry et al., 1990; Papaioannou and Behringer, 2005; Yang et al., 2010; Nandi and Mishra, 2015).

**Supplementary Table 3**. Presentation of data used for generating the dot plots with color scaling in Fig. 4D, D'. Embryos collected from control and CBZ-exposed females at GD9.5 are aligned based on their number of somites (column 1) together with their total scoring (columns 3,4). Number of embryos in each group (column 2) was used for generating the different dot sizes. Female's age in weeks is shown in column 7.

| Number of somites | # of embryos | Total scoring | Average scoring | Female age (weeks) | Group |
| --- | --- | --- | --- | --- | --- |
| 19 | 2 | 13 | 13.5 | 8 | Control |
| 19 |  | 14 |  | 8 | Control |
| 20 | 1 | 15 | 15 | 8 | Control |
| 22 | 1 | 15 | 15 | 8 | Control |
| 23 | 2 | 16 | 16.5 | 8 | Control |
| 23 |  | 17 |  | 8 | Control |
| 24 | 8 | 14 | 15.75 | 8 | Control |
| 24 |  | 15 |  | 8 | Control |
| 24 |  | 15 |  | 8 | Control |
| 24 |  | 16 |  | 8 | Control |
| 24 |  | 18 |  | 8 | Control |
| 24 |  | 15 |  | 8 | Control |
| 24 |  | 16 |  | 8 | Control |
| 24 |  | 17 |  | 8 | Control |
| 25 | 6 | 16 | 16.2 | 8 | Control |
| 25 |  | 17 |  | 8 | Control |
| 25 |  | 15 |  | 8 | Control |
| 25 |  | 15 |  | 8 | Control |
| 25 |  | 16 |  | 8 | Control |
| 25 |  | 18 |  | 8 | Control |
| 26 | 2 | 15 | 16.5 | 8 | Control |
| 26 |  | 18 |  | 8 | Control |
| 27 | 2 | 16 | 15.5 | 8 | Control |
| 27 |  | 15 |  | 8 | Control |
| 28 | 1 | 15 | 15 | 8 | Control |
| 29 | 2 | 17 | 17 | 8 | Control |
| 29 |  | 17 |  | 8 | Control |
| 15 | 1 | 9 | 9 | 9 | Control |
| 20 | 1 | 14 | 14 | 9 | Control |
| 23 | 1 | 16 | 16 | 9 | Control |
| 25 | 2 | 17 | 16.5 | 9 | Control |
| 25 |  | 16 |  | 9 | Control |
| 26 | 1 | 18 | 18 | 9 | Control |
| 13 | 1 | 6 | 6 | 10 | Control |
| 23 | 1 | 15 | 15 | 10 | Control |
| 25 | 1 | 16 | 16 | 10 | Control |
| 27 | 2 | 18 | 18.5 | 10 | Control |
| 27 |  | 19 |  | 10 | Control |
| 29 | 1 | 19 | 19 | 10 | Control |
| 30 | 1 | 19 | 19 | 10 | Control |
| 31 | 3 | 19 | 19 | 10 | Control |
| 31 |  | 19 |  | 10 | Control |
| 31 |  | 19 |  | 10 | Control |
| 32 | 1 | 19 | 19 | 10 | Control |
| 34 | 1 | 20 | 20 | 10 | Control |
| 35 | 1 | 19 | 19 | 10 | Control |
| 14 | 1 | 7 | 7 | 11 | Control |
| 16 | 1 | 12 | 12 | 11 | Control |
| 17 | 1 | 12 | 12 | 11 | Control |
| 18 | 1 | 12 | 12 | 11 | Control |
| 19 | 1 | 13 | 13 | 11 | Control |
| 20 | 3 | 14 | 14.3 | 11 | Control |
| 20 |  | 16 |  | 11 | Control |
| 20 |  | 13 |  | 11 | Control |
| 22 | 1 | 15 | 15 | 11 | Control |
| 24 | 1 | 17 | 17 | 11 | Control |
| 27 | 1 | 16 | 16 | 11 | Control |
| 29 | 1 | 16 | 16 | 11 | Control |
| 4 | 1 | 5 | 5 | 14 | Control |
| 14 | 1 | 13 | 13 | 14 | Control |
| 16 | 1 | 13 | 13 | 14 | Control |
| 19 | 2 | 14 | 14 | 14 | Control |
| 19 |  | 14 |  | 14 | Control |
| 22 | 1 | 17 | 17 | 14 | Control |
| 23 | 1 | 17 | 17 | 14 | Control |
| 24 | 2 | 16 | 16.5 | 14 | Control |
| 24 |  | 17 |  | 14 | Control |
| 25 | 2 | 17 | 17 | 14 | Control |
| 25 |  | 17 |  | 14 | Control |
| 26 | 3 | 18 | 17.7 | 14 | Control |
| 26 |  | 18 |  | 14 | Control |
| 26 |  | 17 |  | 14 | Control |
| 20 | 5 | 17 | 15.4 | 8 | CBZ |
| 20 |  | 14 |  | 8 | CBZ |
| 20 |  | 15 |  | 8 | CBZ |
| 20 |  | 16 |  | 8 | CBZ |
| 20 |  | 15 |  | 8 | CBZ |
| 21 | 1 | 16 | 16 | 8 | CBZ |
| 22 | 3 | 17 | 16.3 | 8 | CBZ |
| 22 |  | 15 |  | 8 | CBZ |
| 22 |  | 17 |  | 8 | CBZ |
| 23 | 1 | 17 | 17 | 8 | CBZ |
| 24 | 2 | 17 | 17 | 8 | CBZ |
| 24 |  | 17 |  | 8 | CBZ |
| 25 | 4 | 18 | 17 | 8 | CBZ |
| 25 |  | 17 |  | 8 | CBZ |
| 25 |  | 16 |  | 8 | CBZ |
| 25 |  | 17 |  | 8 | CBZ |
| 26 | 1 | 18 | 18 | 8 | CBZ |
| 27 | 1 | 18 | 18 | 8 | CBZ |
| 2 | 2 | 5 | 5 | 10 | CBZ |
| 2 |  | 5 |  | 10 | CBZ |
| 18 | 1 | 14 | 14 | 10 | CBZ |
| 19 | 1 | 15 | 15 | 10 | CBZ |
| 19 | 1 | 14 | 14 | 10 | CBZ |
| 20 | 1 | 16 | 16 | 10 | CBZ |
| 21 | 1 | 13 | 13 | 10 | CBZ |
| 15 | 1 | 10 | 10 | 11 | CBZ |
| 18 | 1 | 14 | 14 | 11 | CBZ |
| 19 | 1 | 14 | 14 | 11 | CBZ |
| 20 | 3 | 15 | 14.7 | 11 | CBZ |
| 20 |  | 14 |  | 11 | CBZ |
| 20 |  | 15 |  | 11 | CBZ |
| 21 | 2 | 15 | 14.5 | 11 | CBZ |
| 21 |  | 14 |  | 11 | CBZ |
| 22 | 3 | 16 | 15 | 11 | CBZ |
| 22 |  | 14 |  | 11 | CBZ |
| 22 |  | 15 |  | 11 | CBZ |
| 23 | 2 | 16 | 16 | 11 | CBZ |
| 23 |  | 16 |  | 11 | CBZ |
| 24 | 3 | 14 | 15.3 | 11 | CBZ |
| 24 |  | 16 |  | 11 | CBZ |
| 24 |  | 16 |  | 11 | CBZ |
| 25 | 1 | 15 | 15 | 11 | CBZ |
| 26 | 1 | 17 | 17 | 11 | CBZ |
| 14 | 2 | 10 | 11 | 12 | CBZ |
| 14 |  | 12 |  | 12 | CBZ |
| 16 | 1 | 11 | 11 | 12 | CBZ |
| 18 | 1 | 15 | 15 | 12 | CBZ |
| 19 | 3 | 14 | 14.7 | 12 | CBZ |
| 19 |  | 15 |  | 12 | CBZ |
| 19 |  | 15 |  | 12 | CBZ |
| 21 | 3 | 14 | 14.7 | 18 | CBZ |
| 21 |  | 15 |  | 18 | CBZ |
| 21 |  | 15 |  | 18 | CBZ |
| 23 | 4 | 18 | 16.3 | 18 | CBZ |
| 23 |  | 17 |  | 18 | CBZ |
| 23 |  | 16 |  | 18 | CBZ |
| 23 |  | 14 |  | 18 | CBZ |
| 24 | 1 | 18 | 18 | 18 | CBZ |
| 25 | 1 | 18 | 18 | 18 | CBZ |
| 27 | 1 | 19 | 19 | 18 | CBZ |
| 18 | 1 | 13 | 13 | 19 | CBZ |
| 19 | 1 | 13 | 13 | 19 | CBZ |
| 20 | 3 | 13 | 13.7 | 19 | CBZ |
| 20 |  | 14 |  | 19 | CBZ |
| 20 |  | 14 |  | 19 | CBZ |
| 21 | 1 | 14 | 14 | 19 | CBZ |
| 23 | 3 | 15 | 15 | 19 | CBZ |
| 23 |  | 15 |  | 19 | CBZ |
| 23 |  | 15 |  | 19 | CBZ |
| 24 | 1 | 13 | 13 | 19 | CBZ |
| 25 | 1 | 16 | 16 | 19 | CBZ |
| 27 | 1 | 16 | 16 | 19 | CBZ |
| 28 | 1 | 19 | 19 | 19 | CBZ |
| 29 | 1 | 18 | 18 | 19 | CBZ |
| 31 | 1 | 19 | 19 | 19 | CBZ |

**Supplementary Figure 1**. The effect of CBZ on apoptotic cell death

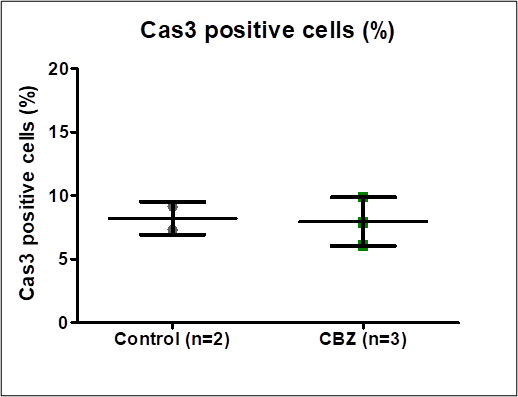

Control or CBZ-exposed embryos were immune-stained for Caspase 3 (Cas3) and analyzed by flow-cytometry to test for apoptotic cells. Percentage of Cas3-expressing cells out of the total cells is shown for each group, with no significant differences between the groups (p=0.3). Each dot represents total cells from a pool of 2-3 different embryos from 2 control females and 3 CBZ-exposed females.
